# Effects of Sequence Composition and Patterning on the Structure and Dynamics of Intrinsically Disordered Proteins

**DOI:** 10.1101/2020.06.08.137752

**Authors:** Andrei Vovk, Anton Zilman

## Abstract

Unlike the well defined structures of classical natively folded proteins, Intrinsically Disordered Proteins (IDP) and Intrinsically Disordered Regions (IDR) dynamically span large conformational and structural ensembles. This dynamic disorder impedes the study of the relationship between the amino acid sequences of the IDPs and their spatial structures, dynamics, and function. Multiple experimental and theoretical evidence points in many cases to the overall importance of the general properties of the amino acid sequence of the IPDs rather than their precise atomistic details. However, while different experimental techniques can probe aspects of the IDP conformations, often different techniques or conditions offer seemingly contradictory results. Using coarse-grained polymer models informed by experimental observations, we investigate the effects of several key variables on the dimensions and the dynamics of IDPs. The coarse-grained simulations are in a good agreement with the results of atomistic MD. We show that the sequence composition and patterning are well reflected in the global conformational variables such as the radius of gyration and hydrodynamic radius, while the end-to-end distance and dynamics are highly sequence specific. We identify the conditions that allow mapping of highly heterogeneous sequences of IDPs onto averaged minimal polymer models. We discuss the implications of these results for the interpretation of the recent experimental measurements, and for further development of appropriate mesoscopic models of IDPs.

## 1 Introduction

Many proteins are intrinsically disordered and do not conform to the classical structure - function paradigm. Yet, these proteins possess diverse biological functions, while maintaining high dynamic and structural flexibility. Under native conditions, their structures comprise dynamic ensembles of different conformations. Intrinsically disordered proteins (IDPs) or intrinsically disordered regions (IDRs) became the common nomenclature used to distinguish this class of proteins and peptides from the traditional ordered proteins [1, 2]. IDPs are involved in a wide range of health and disease processes and functions of the cell. Furthermore, a broad array of human diseases are associated with the failure of an ordered protein to adopt its native conformation, consequently gaining some of the properties of an IDP [2, 3]. Proteins associated with cancer, diabetes, neurodegenerative, and cardiovascular diseases often have regions of structural disorder, making them the leading targets for drug development [1, 2, 4].

Understanding how an IDP’s amino acid sequence dictates the equilibrium and the dynamical properties of its conformational ensemble is an important step towards understanding the principles of function of this class of proteins. A full characterization of an IDP, in principle, involves a description of all possible conformational states and the rates of inter-conversion between them, which is hard to access experimentally [5]. Nevertheless, several experimental techniques reveal information about various characteristics of the IDP ensembles: NMR, fluorescence correlation spectroscopy (FCS), or dynamic light scattering (DLS) can measure the diffusion coefficient and the corresponding hydrodynamic radius of an IDP, Förster resonance energy transfer (FRET) provides information about the inter-residue distances (such as the end-to-end distance), and small angle X-ray scattering (SAXS) can measure the radius of gyration [2, 6, 7].

Emerging evidence shows that due to their disordered nature and the importance of entropic effects, IDP structural ensembles might be less sensitive to the fine details of a specific amino acid sequence compared to the unique 3D structures of the classical folded proteins. Rather, many IDP properties can often be understood in terms of the global characteristics such as the overall charge, hydrophobicity, flexibility of the polypeptide backbone, and the average solvent properties [8]. Typically, mean hydrophobicity is lower and the mean net charge is higher in IDP sequences than in folded proteins, and they are impoverished in large amino acids, preventing the folding of IDPs into stable unique structures with a hydrophobic core. A predictor based only on the amino acid composition predicts disorder with 87% accuracy [9]. A predictor based on the reduction of the size of the sequence alphabet by assigning each amino acid to just one of 4 types (neutral, hydropbobic, positive and negative), performs almost as well as a predictor using the full 20 amino acid alphabet [9]. Even a minimal predictor based only on two properties: the net charge per residue and and the mean hydrophobicity per residue, can often differentiate well between IDPs and folded proteins, as well as between different classes of IDPs [1–3, 6, 10, 11].

Polymer physics offers a useful theoretical framework for understanding IDP behaviors, and enables linking experimental observables to the underlying conformational ensembles [2, 12, 13]. Simple mean field homopolymer models have been successful in categorizing the IDP ensembles into regimes of qualitatively different behaviours based on the ensemble averages of polymer dimensions, such as the radius of gyration and the end-to-end distance [12–14]. Commonly, the size of an IDP chain in space correlates with the net balance between the repulsive and attractive intra-chain and chain-solvent interactions, which can often be encapsulated in an effective internal cohesiveness parameter, related to the classical Flory parameter *χ* [11, 15–18]. The ratio of the fraction of the charged amino acids to the fraction of hydrophobic ones is often sufficient to distinguish between the swollen and compact regimes of behaviour [11, 19, 20].

At the low cohesiveness, disordered, extreme polypeptides are often successfully described by models of polymers in a good solvent and adopt diffuse swollen random coil conformations. In the opposite, high cohesiveness, regime the IDPs adopt dense globular conformations [12, 16, 21]. In particluar, IDP location on the order-disorder continuum can often be encapsulated in the scaling dependence of their size *R* on the chain length (number of amino acids) *N*, *R* ~ *N^ν^*, which describe the universal features of the behaviour of polymeric molecules that are largely independent of the details of the local microscopic properties of the chain or the solvent [1, 12, 14, 22–25]. In the highly disordered regime (such as at high denaturant concentrations and low intra-chain cohesiveness), the IDP dimensions follow the good solvent scaling law *ν* ≃ 0.6, which gradually decreases to *ν* ≃ 1/3 in the compact globular regime at high cohesiveness. These simple mean field theories have been successful not only in describing individual molecules of IDPs but also multi-chain systems in various geometries - from surface grafted layers to 3D phase separation [17, 18, 26–28].

However, despite their successes, simple mean field polymer theories suffer from several drawbacks. First, they fail to differentiate between distinct polymer dimensions, such as the end-to-end distance, radius of gyration, and the hydrodynamic radius, which can lead to difficulties in the interpretation of the experimental data. Several recent works using FRET and SAXS measurements unveiled discrepancies and divergent behaviors of the different measures of polymer dimensions [29–33]. In particular, the chain radius of gyration *R_g_*, inferred from FRET measurements of the end-to-end distance *R_e_* can show much greater compaction with the decrease in the denaturant concentration compared to the direct SAXS measurement of *R_g_* [29]. Similar “decoupling” between the *R_g_* and the end-to-end distance *R_e_* was observed in [33]. On the other hand [31] observed consistent increase in all chain dimension with an increase in the denaturant concentration, using multiple methods: FRET for *R_e_*, SAXS for *R_g_*, and FCS and DLS for the hydrodynamic radius *R_h_.* One proposed explanation for such decoupling is the effect of FRET dyes located at the chain ends [30, 34, 35]. On the other hand, [32] and [33] did not report an observable effect of the dyes on the chain dimensions. These results raise important and fundamental questions about the methodology of inference of the chain dimensions and internal structures of IDPs from the experimental data, which may depend on the specific assumptions in the polymer models used [32, 33].

Second, simple polymer theories fail to capture the effects of sequence heterogeneity. Although some atomistic details may be coarse-grained [6, 13, 36–38], the effects and the importance of the amino acid patterning on the dimensions and the dynamics of IDPs are not fully known [11, 12, 14, 39, 40]. In particular, permutations of the sequences of the amino acids without changing the overall composition may affect the polymer dimensions as predicted computationally [14, 39, 40] and observed experimentally [25]. Similarly, as mentioned above, specific amino acids located near the ends of the chain might have strong effects on some of the chain properties. Furthermore, hitherto not fully explained inconsistencies arise in the measurements of the dynamic reconfiguration times of the IDPs, explored via FRET and Fluorescence Correlation Spectroscopy (FCS) [7, 41–43].

Interpretation of the experimental data often relies on the computational models of IDPs. As mentioned above, simple mean field polymer models are powerful but not sufficient to capture the complexity of the whole gamut of behaviors of IDPs. Computational approaches based on computer simulations offer a way to systematically study the vast sequence space and the effects of sequence heterogeneity on the polymer dimensions and other properties. On the one hand, all atom molecular dynamics (MD) simulations have been used as a tool in the modeling of natively folded proteins for several decades. However, there are several obstacles when applying these methods to IDPs. Even with the dramatic increases in the computing power, computationally expensive simulations required to fully explore the vast conformational space of an IDP are not always feasible [5, 44]. Moreover, agreed upon atomistic force fields for IDPs are still lacking, and their predictions remain sensitive to the fine tuned choices of parameter values, and are potentially prone to over-fitting [45–48].

On the other hand, coarse-grained simulations avoid many of these pitfalls by subsuming many atomistic details into the coarse grained variables, such as the local amino acid charge, hydrophobicity and monomer size [49–55]. Identification of the key properties and molecular features that capture the connection between the IDP structure and the experimentally accessible variables [50, 51] while avoiding over-fitting the sparse experimental data is challenging [51]. Several of these properties have been identified: the importance of electrostatic interactions, hydrophobicity and, more generally, the association of certain amino acids with either expansion or compaction of the IDPs. Yet, although a number of different force fields and solvent models have been successfully applied in different specific cases, currently there are no universally accepted coarse-grained (or atomistic) force fields. In order to reproduce the experimental data, simulation outcomes often require sub-ensemble sampling and re-weighting [31, 33, 56], or additional *ad hoc* assumption about the ensemble properties [7, 29, 31, 33, 41, 56]. Furthermore, the link between atomistic and coarse-grained models is still missing, as is the understanding of the regimes of applicability of different approaches.

Identification of quantitative characteristics of the IDP sequence that enable the link between the molecular properties and the macroscopic behavior is desirable for further progress. This is of particular importance for translating the insights of studies of individual IDP molecules towards understanding and prediction of their collective properties, such as phase separation and coacervation [57, 58]. In this paper we systematically investigate the effects of sequence composition and heterogeneity on the dimensions and the dynamics of IDP conformational ensembles. We use experiment-informed coarsegrained minimal complexity models that include only the key features of an IDP sequence. We specifically focus on the effects of the chain “patchiness” and the sequence near the chain ends. We investigate the mapping between the atomistic and coarse-grained descriptions, and identify coarse grained parameters and variables that enable mapping from heterogeneous onto averaged models. The results shed light on the interpretation of recent experimental results and serve as a basis for further development of mesoscopic models of IDPs.

The paper is structured as follows. In Section 2, we describe the computational methods of the paper based on the over-damped Langevin dynamics with explicit hydrodynamic interactions. In Section 3.1.1, we present the results of the simulations of a minimal homopolymer model of intra-chain interactions to differentiate between the various polymer dimensions: end-to-end distance, radius of gyration and hydrodynamic radius. In Section 3.1.2, we investigate the effects of sequence heterogeneity on the IDP dimensions expanding the homopolymer model to include four monomer types (cohesive, neutral, positively charged, or negatively charged). In Section 3.2, we study the effects of the amino acid sequence on the end-to-end dynamics of IDPs and discuss the implication for interpretation of experimental results. We conclude with Conclusions and Discussion in Section 4.

## 2 Methods

We represent an IDP as a polymer consisting of *N* monomers. Sequence effects are introduced by assigning each monomer into one of the four types: neutral, cohesive, positively charged, or negatively charged. Similar models and computational implementations have been used to study polymer systems [49, 59–63]. The model can accommodate various levels of detail such as sequence heterogeneity and hydrodynamic interactions.

The monomers are kept on a chain via the finitely extensible non-linear elastic potential (FENE) bonds between nearest neighbor monomers [64]:

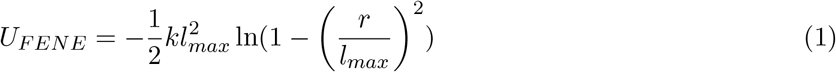

All monomer pairs interact via a repulsive 8-6 LJ potential modeling the steric repulsion between the monomers:

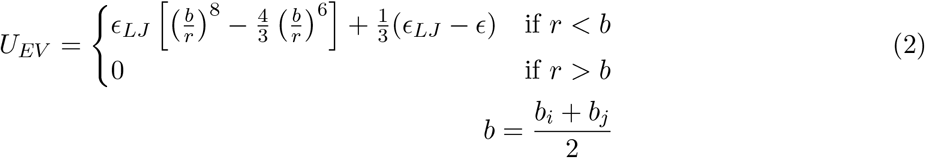

where *ϵ_LJ_* is the strength of the repulsion, and *b* (equal to the sum of the radii of the two interacting monomers) is the distance between the monomer centers where the force is zero. An exception to this rule occurs if the two interacting beads are bonded monomers of a polymer: in this case *b* = *b*_0_, which reflects the bond length rather than the radius. The potential is shifted by 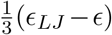 in order to maintain continuity at *r* = *b* with the attractive potential described in the following paragraph.

In addition to the universal repulsive interaction, “cohesive” monomers interact through the attractive potential

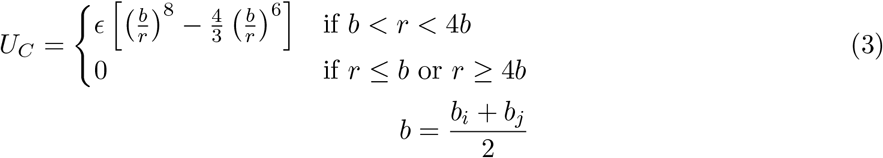

The parameter *ϵ* controls the strength of the attraction between the monomers. The sum of the radii of the two beads (*b*) is the same as in the repulsive force described previously. The attractive potential smoothly splines with the repulsive part at *r* = *b*. To reduce computational complexity, the potential is cut off beyond *r* = 4*b*, where it is ~ 0.1% of it’s maximal depth.

Interaction between two charged monomers is modeled via the screened Coulomb potential:

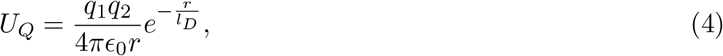

where *q*_1_ and *q*_2_ are the charges of the beads, and *ϵ*_0_ is the dielectric permittivity of the solution. The Debye length *l_D_* describes the screening of the electrostatic potential by the salt ions.

The dynamics of the chain are described by the over-damped Langevin dynamics implemented via Ermak-McCammon [65] algorithm, as described below. Hydrodynamic interactions are included via the Rotne-Prager-Yamakawa tensor [66, 67]. T

For convenience, we define the following dimensionless variables: the position of a monomer 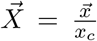, the simulation time step 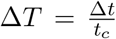, the sum of the deterministic forces on a monomer due to it’s interactions with the other monomers 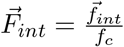. The units of force are 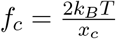, the units of length are 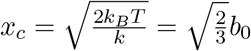 (*k* and *b*_0_ are defined above), and the units of time are 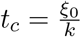, where *ξ*_0_ = 6*πηa*_0_, is the Stokes drag coefficient for a bead with hydrodynamic radius *a*_0_. In these units, the displacement of a monomer in one simulation time step is:

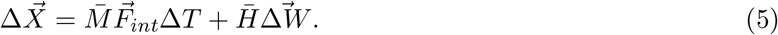

When hydrodynamic interactions are included, 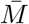 is the Rotne-Prager-Yamakawa tensor [67] multiplied by *ξ*_0_, and 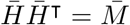. In simulations, 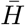 is chosen to be a lower triangular matrix obtained using the Cholesky decomposition of 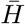. For the calculation of the equilibrium quantities, such as the radius of gyration of the end-to-end distance, hydrodynamic interactions are immaterial, and all off-diagonal entries of 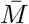 may be set to 0. The components Δ*W_i_* of 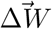 are independent random variables with Gaussian distributions such that 〈Δ*W_i_*〉 = 0 and 〈Δ*W_i_*(*T*)Δ*W_j_*(*T*′)〉 = Δ*Tδ*(*T*′ – T)*δ_ij_* [65, 68, 69]; see Appendix for details.

Expressed in the simulation units, the range (diameters) of the repulsive volume interactions between bonded monomers is 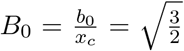 The maximal extension of the FENE bonds between monomers is 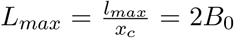. The strength of excluded volume interactions is 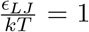. The hydrodynamic radii of the monomers were 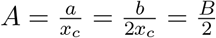.

## 3 Results and Discussion

### 3.1 Effects of sequence and interactions on the chain dimensions

#### 3.1.1 Effects of internal cohesiveness on the chain dimensions: averaged homopolymer models

Several recent experiments reported discrepancies between polymer dimensions of IDPs/chemically denatured proteins measured using different experimental techniques, most prominently FRET and SAXS [29, 30]. Many of these discrepancies may result from different a choices of the polymer model, of the force field and the re-sampling procedure [31, 33, 56].

In this section, we explore the effects of the intra-chain interactions on the polymer conformational ensemble, and the corresponding experimentally relevant dimensions, such as the end-to-end distance *R_e_*, radius of gyration *R_g_* and the hydrodynamic radius *R_h_.*

These dimensions are defined as:

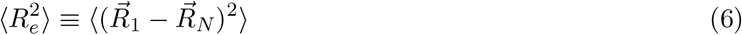

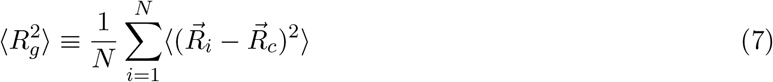

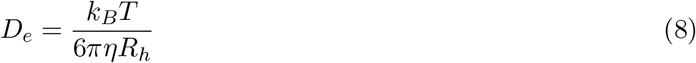

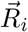 is the position of monomer *i* and 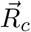 is the polymer centre of mass. *D_e_* is the diffusion coefficient of the polymer centre of mass. The Kirkwood approximation for the hydrodynamic radius is (see Appendix A.1):

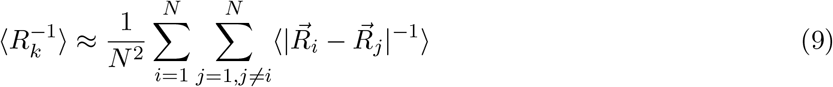

In this section we use a minimal homopolymer model which serves as a “null hypothesis” for the interpretation and the analysis of the experimental data, against which more complex models can be benchmarked. In the model, all monomers of the chain interact attractively with each other with the same average interaction strength *ϵ* (see Equation 3). This coarse-grained interaction parameter subsumes all the direct and solvent-mediated interactions between the monomers properties of the solvent, as well as the average composition and the sequence details of an IDP. Experimentally, low *ϵ* ≃ 0 represents a protein in a high denaturant conditions or an IDP with many disorder promoting amino acids in its sequence. Increasing *ϵ* represents lower denaturant concentration or higher fraction of order promoting amino acids in an IDP sequence.

The cohesiveness parameter *ϵ* is closely related to the classical mean field Flory interaction parameter *χ* [15], which encapsulates all the information about an IDP’s sequence and molecular properties; mathematically the two are related through the second virial coefficient of the interaction *χ* ≃ ∫ *d*^3^*r*(1 – *e*^−*U*(*r*)^, where *U*(*r*) is defined in equations 3 and 2. Unlike mean-field models, the simulations are able to differentiate between the various polymer dimensions: end-to-end distance, radius of gyration and hydrodynamic radius.

Simulations were performed for chains of *N* = 100 monomers and cohesive interactions strengths ranging from 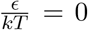 to 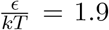 inclusive, in intervals of 0.1. For each *ϵ*, 40 independent runs were performed, each lasting 10^8^ steps, with a time step of Δ*T* = 0.001. Each run began from a self-avoiding random walk initial condition. The first 10^6^ steps were excluded from the analysis in order to avoid biasing the results by the initial conditions, and averages were taken over the time steps and the different runs.

The results are summarized in Figure 1*a*, which shows the average end-to-end distance, the radius of gyration, and the hydrodynamic radius. The end-to-end distance has been scaled down by a factor of 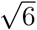 to be comparable to the other dimensions. Overall, all the polymer dimensions monotonically decrease with increasing *ϵ*, as the chain compacts from a coil to a globule. The *θ*-point, where the inter-monomer repulsion is balanced by the inter-monomer attraction resulting in roughly ideal chain behavior, is located around 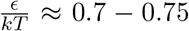 (see Appendix A.2); however the exact location of the *θ*-point may depend on the specific choice of the form of the interaction potential [70]. The end-to-end distance undergoes the greatest relative compaction, while the hydrodynamic radius experiences the least change.

**Figure 1:**
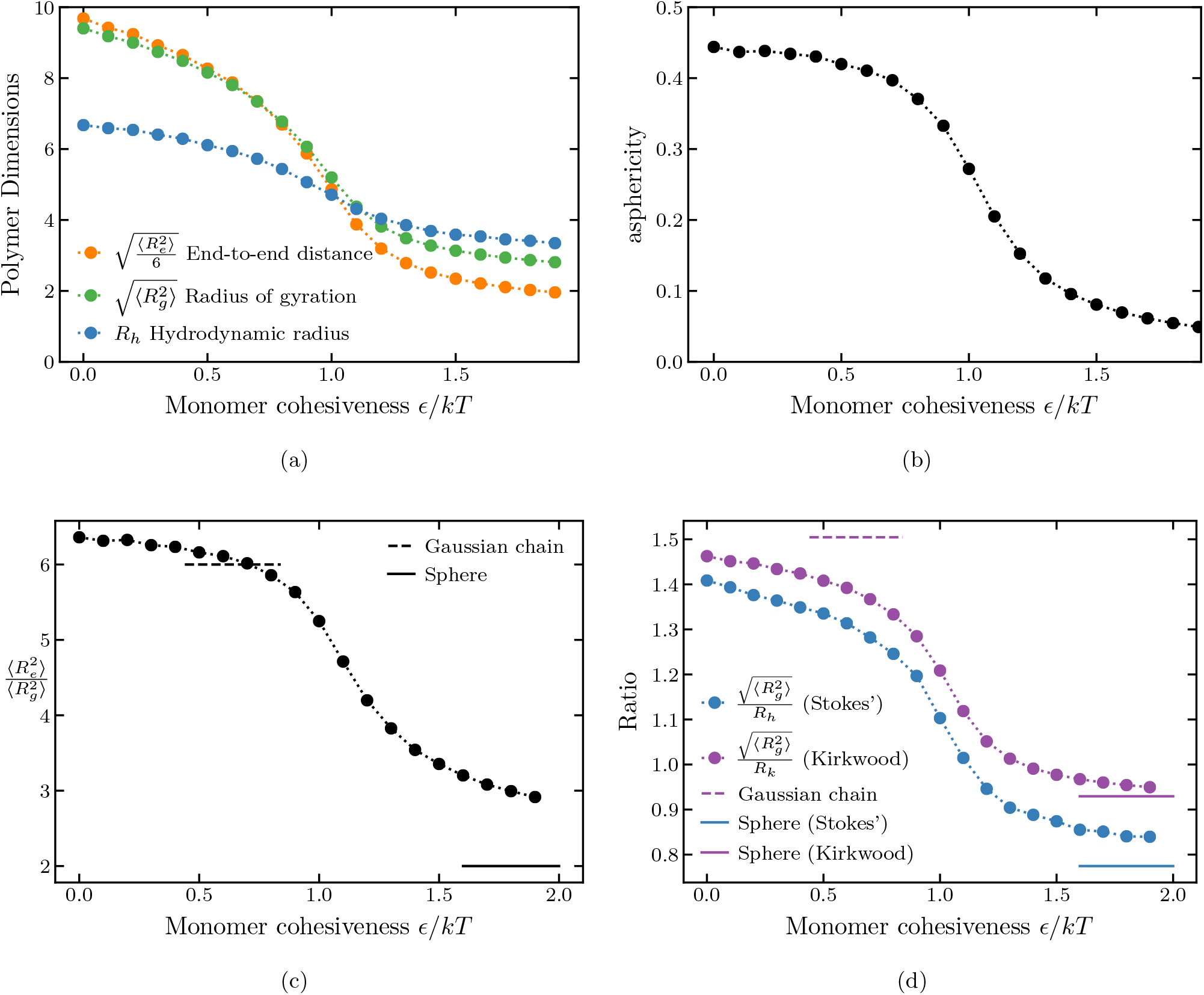
*(a)* Polymer dimensions of a homopolymer for varying monomer cohesiveness. *(b)* Asphericity of a homopolymer for varying monomer cohesiveness. *(c)* Ratio of the square of the end-to-end distance to the square of the radius of gyration of a homopolymer for varying monomer cohesiveness. Solid horizontal line corresponds to a uniform sphere. Dashed horizontal line marks the *θ*-point where the polymer behaves as a Gaussian chain with 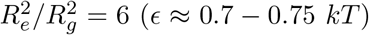. *(d)* Blue: ratio of the radius of gyration to the hydrodynamic radius. Purple: ratio of the radius of gyration to the Kirkwood approximation to the hydrodynamic radius. The good solvent corresponds to *ϵ* = 0, the *θ*-solvent corresponds to *ϵ* ≈ 0.7 — 0.75 *kT*, and poor solvents correspond to *ϵ* > 1.5 *kT*. The number of monomers is *N* = 100.

One can more readily identify a polymer position on the order-disorder continuum by studying the ratios between the various polymer dimensions rather than the individual dimensions themselves in isolation. As will be seen in the next section, these ratios can be more robust and versatile measures of the polymer conformations.

Figures 1*c* and 1*d* show the ratios of the square of the end-to-end distance to the square of the radius of gyration, as well as the ratio of the radius of gyration to the hydrodynamic radius for varying values of polymer cohesiveness *ϵ*. The ratios obtained from simulations approach the theoretical limits for good, θ, and poor solvents (calculated for *N* → ∞). For the self-avoiding walk (*ϵ* = 0), 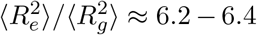 [71, 72](depending on the approximation). For an ideal chain (*θ* point), 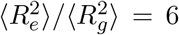. In the compact regime of high cohesiveness, the polymer can be approximated as a uniformly dense sphere. In this regime, assuming that the locations of the two ends are independent from each other and are uniformly distributed inside the sphere, 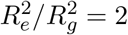 [56, 73]. The ratio of the radius of gyration to the hydrodynamic radius is known to be *R_g_*/*R_k_* ~ 1.5 for the *θ* solvent [23, 74] and decreases to *R_g_*/*R_h_* ~ 0.774 and *R_g_*/*R_k_* ~ 0.93 in the globular high cohesiveness regime [56, 73]. Importantly, in the homopolymer model the *R_g_* and *R_e_* remain coupled in a sense that both consistently decrease with the increase in *ϵ*.

##### Chain asphericity

As mentioned above, some of the discrepancies between FRET and SAXS measurements of the radii of gyration can be attributed to the assumptions of the hompolymer models used in the inference of polymer dimensions from the data. In particular, asphericity (sometimes referred to as the shape anisotropy) *δ* of IDP ensembles has been proposed to play an important role in the inference of IDP properties form FRET and SAXS data [33, 56]. Although the ensemble average monomer density is isotropic for any polymer, the individual conformations may not be, giving a non-zero average asphericity. For a rigid rod *δ* = 1, and for a sphere *δ* = 0. The ensemble averaged asphericity is:

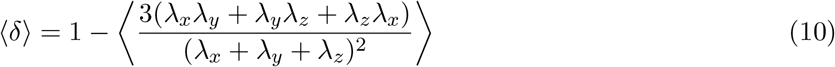

where *λ_x_*, *λ_y_*, and *λ_z_* are the eigenvalues of the 3 × 3 gyration tensor for a single conformation, whose entries are:

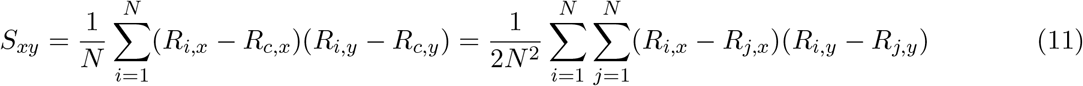

*R_i,x_* and *R_c,x_* are the *x*-components of the position of monomer *i* and the center of mass respectively. The radius of gyration for that conformation is: 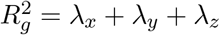.

Figure 1*b* shows the asphericity of a homopolymer chain for different values of monomer cohesiveness and decreases from ~ 0.45 for the swollen coil to close to 0 for the compact globular conformations. For the homopolymer model, the asphericity is well correlated with the ratio of the end-to-end distance to the radius of gyration *R_e_*/*R_g_*.

##### Conditional Sub-Ensemble Distributions

Due to the absence of universally accepted force fields to describe the conformational ensembles of the IDPs, sub-ensembles with appropriate conditional distributions of the end-to-end distance conditioned on a sub-ensembles with set values of *R_G_* are commonly used for comparison with the experimental data [29, 56, 75, 76].

In Figure 2, we compare the conditional distributions of the end-to-end distance, *p*(*R_e_*\*R_g_*), obtained from the homompolymer simulations, with the predictions of the common sub-ensemble model, Sanchez-Haran theory [76, 77], which postulates that the end-to-end distance distribution of the conformations conditioned on a particular radius of gyration is the probability distribution of distances between two random points in a sphere of the radius 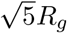.

**Figure 2:**
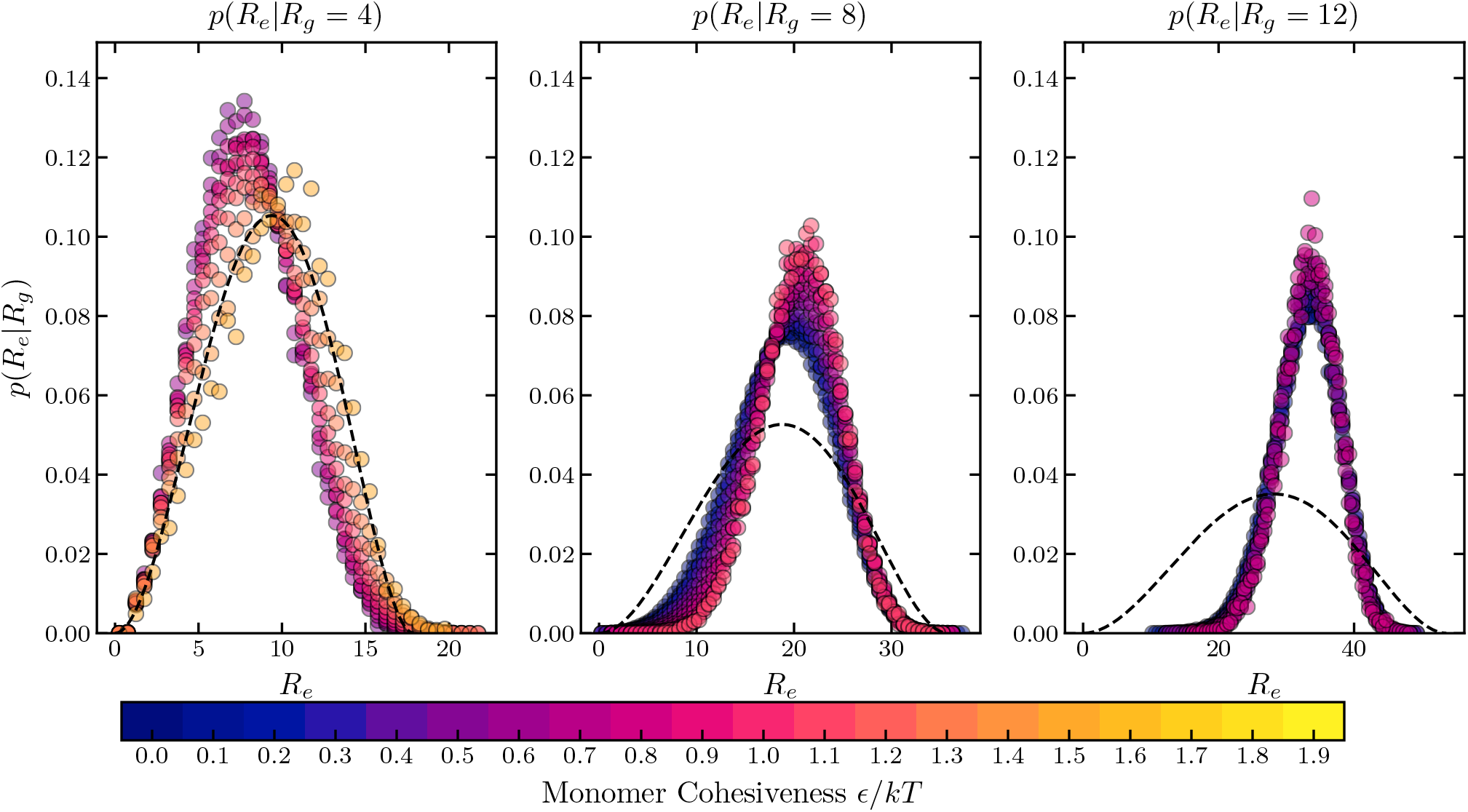
Probability distributions of the end-to-end distance of a homopolymer, conditioned on the sub-ensembles with different radii of gyration. The circle symbols show the simulation results. The color of a symbol (blue to yellow) corresponds to low to high values of *ϵ*. The black dashed line shows the distribution of the end-to-end distance of the Sanchez-Haran model. The number of monomers is *N* = 100. Polymer dimensions are in the units of 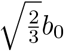 where *b*_0_ is the monomer diameter. Histogram bin size for calculation of the distribution is 0.5; see Section 2.

Notably, the simulated conditional distributions are not noticeably affected by the strength of the cohesive interaction *ϵ*. The Sanchez-Haran distribution matches the simulations well for compact conformations, which typically have large *ϵ*, but underestimates the end-to-end distance for large conformations, which typically have small *ϵ*. Thus, Sanchez-Haran model would tend to overestimate the radius of gyration for polymers with low cohesiveness or in good solvents, based on the raw FRET data.

Another notable artefact of the Sanchez-Haran model is that implicitly assumes that *R_g_*/*R_e_* = 6 (that of a Gaussian chain) for all values or cohesiveness. Following [76]: 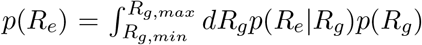. Thus, 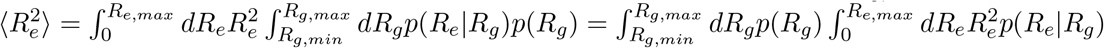. For the distribution of distances between two random points in a sphere of radius 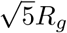, 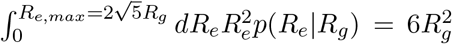 and so this model, like the Gaussian chain model, predicts the relationship 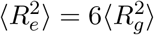.

These results have potentially important implications with respect to the interpretation of FRET and SAXS data.

#### 3.1.2 Effects of Sequence Composition and Patterning

To capture the effects of sequence composition and patterning on IDP structures, we extended the model into the heterogeneous sequence domain. In this section, rather than focusing on specific intrinsically disordered proteins with specific coarse-grained model parameters, we focus on the general relationships between the sequence properties and the polymer dimensions.

As described Section 2, we use a “four letter” model (“HP+-”), where monomers can be either neutral/repulsive (“P”), cohesive/attractive (“H”), positively charged (“+”), or negatively charged (“-”). The first two types of monomers are inspired by the Hydrophobic-Polar model of proteins [78]. The charged monomers represent charged amino acids, while the cohesive monomers can represent order promoting (mostly hydrophobic) amino acids, and the neutral monomers represent polar/disorder promoting amino acids. Overall, this model takes into account the basic features of IDP sequences that typically control their conformations as the polymer dimensions are typically correlated with the compositional balance of the order-promoting and disorder-promoting amino acids [25, 49]. In particular, net charge and hydrophobicity can distinguish IDPs from natively folded proteins [1], and the sequences of charged residues can affect the IDP dimensions dimensions [39].

In the model, neutral monomers experience only repulsive (non-electrostatic) interactions (*ϵ_i_* = 0 and *q_i_* = 0 in Equations 3 and 4). Cohesive monomers interact only with other cohesive monomers via the cohesive interaction (with the strength *ϵ*). Charged monomers interact with other charged monomers via the electrostatic interactions, and via repulsive potentials with non-charged monomers. The bond length between adjacent monomers was 1.35 in simulation units, corresponding to roughly 0.38 nm distance between two adjacent C*α* atoms in real polypeptides. For the sequences comprising mixtures of cohesive (“H”) and neutral monomers (“P”), the steric repulsion diameters of Equation 2 of all monomers were set to 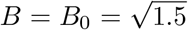 in simulation units, corresponding roughly to ~ 0.35 nm. For the polyampholyte sequences, the steric repulsion diameters were set to *B*_ − = 2.29 and *B*_+_ = 2.44 in simulation units, reflecting the relative volumes of the corresponding amino acids (Lysine “E” and Glutamic acid “K”) [79, 80] (see Section 2). The strength of the electrostatic interactions was *Q* = 2 and the Debye length was *L_D_* = 4 in simulation units corresponding to the screening length of ~ 1.1 nm (typical for ~ 75 mM of NaCl). However, the results apply more generally, and we expect the small variations in the parameterization to not have a major effect on the main results of the paper.

##### Chains composed of neutral and cohesive monomers

We first investigated how the sequence patterning of neutral (“P”) and cohesive (“H”) monomers affects the chain dimensions. We simulated 5 different sequences of 30 cohesive (“H”) and 30 neutral (“P”) monomers using the coarse-grained model. The sequences, shown in Table 1, vary in the sizes of the cohesive and the neutral clusters, increasing from 1 to 5, while maintaining the same 1 : 1 ratio of neutral to cohesive monomers. For each sequence and for each set of interaction parameters *ϵ*, *Q*, and *L_D_*, 8 runs were performed, each lasting 10^8^ steps, with the time step of Δ*T* = 0.001 in simulation units. Each run began with a self-avoiding random walk initial condition. The first 10^6^ steps were excluded from the analysis, and averages were taken over steps over the time and different runs.

**Table 1:**
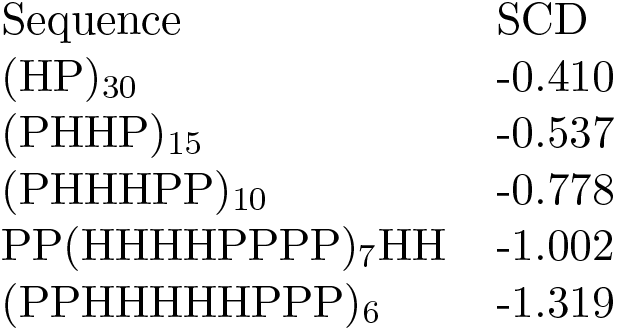
Sequences composed of 30 cohesive (“H”) and 30 neutral (“P”) monomers with different sizes (1, 2, 3, 4, or 5) of cohesive (and neutral) clusters. The subscripts indicate how many times the sequence in parentheses is repeated.

Specifically, we focus on the size of cohesive “patches”, which differs among the sequences while the overall composition stays the same. The “patchiness” of the sequence can be quantified using the Sequence Charge Decoration (SCD) parameter (originally introduced in [81] to describe the patterning of charged monomers). The SCD for the cohesive/neutral sequence is defined in Equation 12,

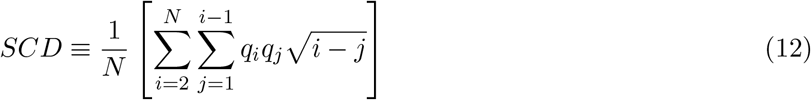

where *N* is the number of the monomers in the sequence, and *q_i_* = +1 for a neutral monomer and *q_i_* = –1 for a cohesive monomers at a position *i*.

The results are summarized in Fig. 3, which explores the effects of the cohesiveness *ϵ* of the “H” monomers and the size of the cohesive “patches” on the polymer dimensions. Results for a corresponding homopolymer of 60 cohesive monomers are shown for comparison. On the *x*-axis, the monomer cohesiveness parameter *ϵ* is rescaled by the square fraction of cohesive monomers.

**Figure 3:**
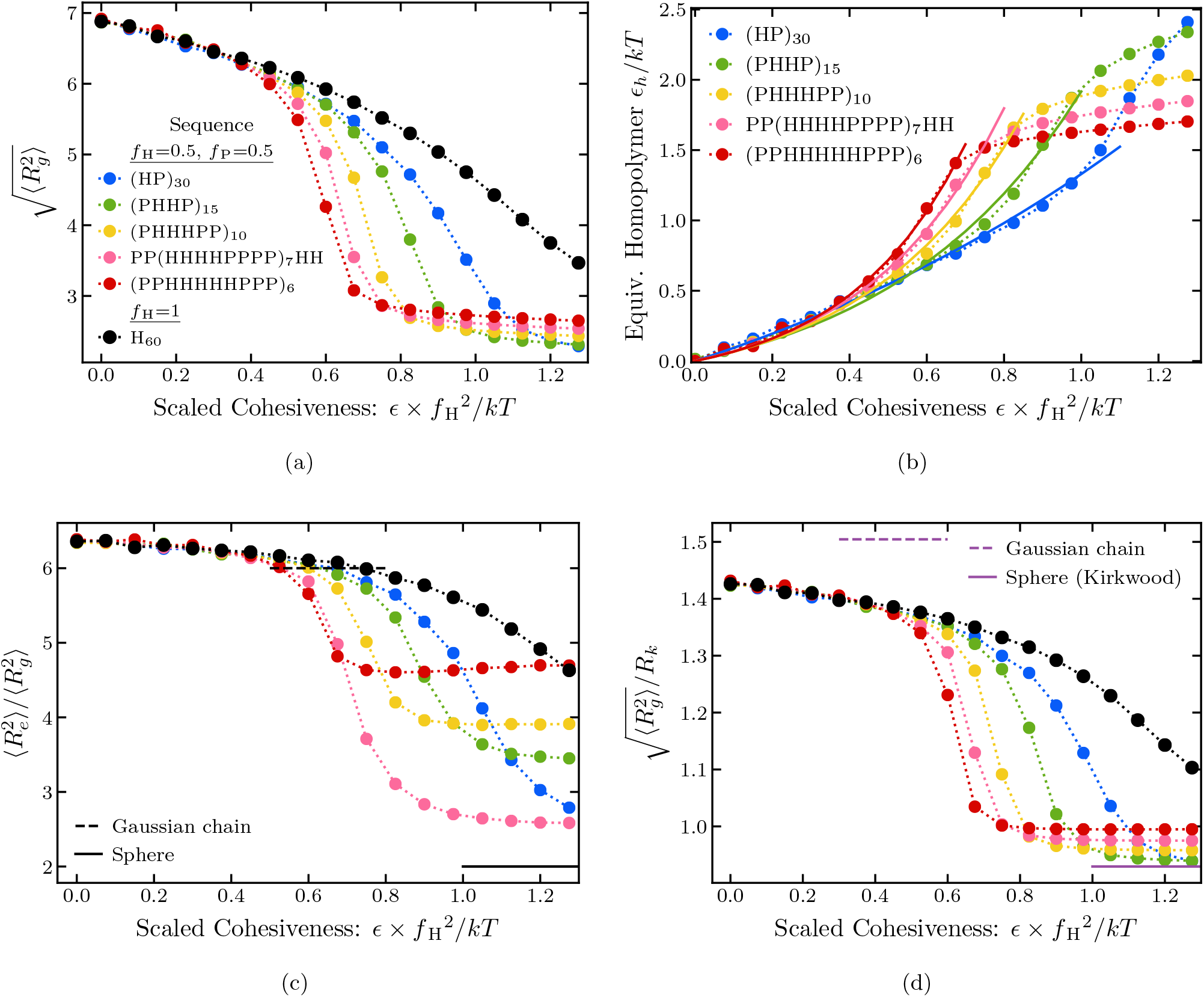
Polymer dimensions as a function of the cohesiveness. *(a)* Radius of gyration for different sequences. *(b)* Dotted line: Equivalent Homopolymer *ϵ*, determined using linear interpolation for 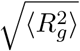. Solid line: empirical fit to (*e^aϵ^* – 1)/b where *a* and *b* are fitting parameters; see text. *(c)* Ratio of the square of the end-to-end distance squared to the square of the radius of gyration. *(d)* Ratio of the radius of gyration to hydrodynamic radius (in Kirkwood approximation). All sequences are composed of 30 cohesive and 30 neutral monomers for varying monomer cohesiveness. The size of the hydrophobic patches varies from 1 to 5; exact sequences are shown in the legend. For comparison, a homopolymer sequence of 60 cohesive monomers is shown in black. The dashed lines correspond to the Gaussian chain predictions, the solid lines correspond to a uniform sphere. *f*_H_ is the fraction of cohesive monomers in the sequence. Radius of gyration is in units of 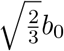 where *b*_0_ is the monomoer diameter, as described in Section 2.

At low cohesiveness, the radii of gyration of all sequences collapse onto an effective homopolymer model with the corresponding value of *ϵ* rescaled by the by the fraction of cohesive monomers squared (*f_H_* = 1/2), reflecting the lower average probability of contacts between cohesive monomers in the heterogeneous sequences. Simple correspondence with the homopolymer begins to break down around the *ϵ* ≈ 0.4 kT. For intermediate cohesiveness, the sequences with larger “patch” sizes exhibit an earlier and steeper coil-to-globule transition. Nevertheless, as shown in Fig. 3a, even moderately cohesive patchy chains can be mapped to an effective homopolymer model with an effective cohesiveness that depends on the size of the cohesive patch (see also Fig. 4).

**Figure 4:**
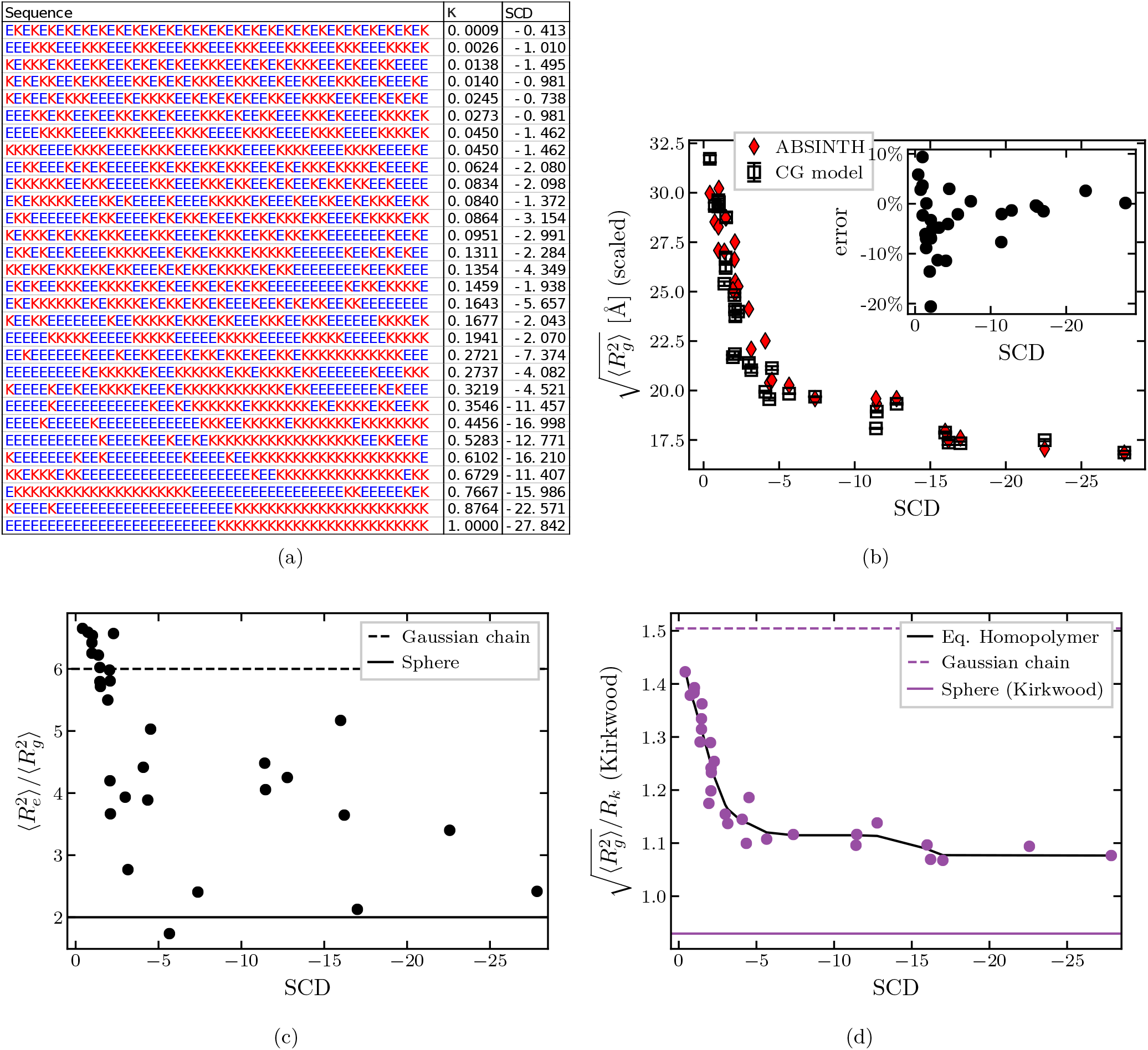
Dimensions of charged polymers. *(a)* Sequences composed of 25 positively and 25 negatively charged amino acids with their corresponding Sequence Charge Decoration (SCD) and *κ* pattern parameters; see text. “K” represents positively charged lysine and “E” represents negatively charged glutamic acid. *(b)* Radii of gyration of the chains with corresponding sequences. Black symbols: coarse-grained model; red symbols: ABSINTH model. *(c)* Squared ratio of the end-to-end distance to the radius of gyration. *(d)* Ratio of the radius of gyration to the hydrodynamic radius (in Kirkwood approximation). Solid black line is the effective homopolymer mapping (see Figure 5). The dashed horizontal lines mark the Gaussian chain predictions, the solid horizonal lines mark the values corresponding to a uniform sphere.

Interestingly, at the high values of cohesiveness in the globular regime the relationship between the the polymer dimensions and the “patch” size is inverted: chains with larger “H” and “P” clusters have larger dimensions. This likely arises from the fact that in this regime “H” “patches” cluster to form a compact cohesive core, decorated by disordered loops of “P” containing spacers.

These trends are reproduced in the behavior of the *R_g_*/*R_h_* ratio, as shown Fig. 3d, and are even more pronounced in the ratio of the end-to-end distance to the radius of gyration (Fig. 3c). These results emphasize that care must be exercised when inferring polymer properties from measurement of polymer dimensions in swollen vs. compact regimes (see more below in Section 7).

We also investigated sequences containing mixtures of cohesive monomers with charges of one type (either positive or negative). Interestingly, the overall results are the very similar to those of the mixtures of cohesive and neutral monomers. Essentially, in this case, charged monomers serve as neutral/repulsive monomers of a renormalized size that is dictated by the Debye length rather than the steric repulsion radius. Full examination of this regime is outside the scope of the present paper, and will be presented elsewhere; see [82].

##### Charged sequences

IDPs commonly contain higher fractions of both positively and negatively charged amino acids in their sequences, compared to the natively folded proteins [12, 14]. In particular, Das and Pappu [39] computationally investigated the effect of charge patterning on the IDP properties using a family of polyampholyte sequences with different degrees of segregation of positive and negative charges in their sequences, shown in Figure 4a. Using Monte Carlo simulations of the IDPs using atomistic ABSINTH force field with implicit solvent [83], they found that the radius of gyration was higher for sequences with well mixed positive and negative charges, and lower for sequences with more segregated charge “patches”. Similar findings were obtained both in the theoretical and the experimental analysis of segregation of order promoting (Proline) and charged residues [25, 40].

In [39], the degree of charge segregation or “patchiness” was quantified using parameter *κ* (defined in the Appendix) whose value is low for well mixed sequences and high for completely segregated sequences. An alternative parameter that quantifies the charge segregation and “patchiness” is known as the Sequence Charge Decoration (SCD) parameter [81], which can be defined for a polyampholyte sequence as in Equation 12 with *q_i_* = 1 for a positively charged monomer and *q_i_* = –1 for a negatively charged one. It has been shown [58] that the radius of gyration simulated by Das and Pappu had a smoother dependence on SCD than on *κ*. Comparison between *κ* and SCD is shown in the Appendix. Other conceptually similar parameters describing the segregation of different types of monomers have been proposed in the literature as well [40]

Figure 4, shows the dependence of the various polymer dimensions on the “patchiness” of the polyampholite sequences (quantified through SCD) calculated using the coarse-grained force field of this paper; see Appendix A.3 for a comparison with *κ* in Figure 9. As shown in Figure 4b, the coarse grained model captures well the overall compaction of the radius of gyration of the chains with the increase the charge “patch” size, as well as the sequence-specific variations in the *R_g_*, compared to the ABSINTH model of [39]. For comparison between our results and those of Das and Pappu [39], our radii of gyration are rescaled by a factor of ~ 1.4 - the ratio of the average radii of gyration over all sequences between our results and those of Das and Pappu. This difference likely arises due to several assumptions of the coarse grained model that differ from the atomistic one: the bond angle restrictions between subsequent amino acids are neglected in the coarse grained model, amino acids are treated as spherically symmetric monomers ignoring the side-chain geometry, and the amino acid size in the LJ steric repulsion potential is based on the volumes of amino acids estimates in folded proteins, which could differ from the excluded volume of amino acids in IDPs [79, 80]. Nevertheless, most of the differences between the two models are below 10 per cent, as shown in the inset of Figure 4b. The sequence with the highest disagreement (of about 20 per cent) is with SCD = 2.070, which comprises repeating periodic motifs of 5 negative amino acids followed by 5 positive ones. This particular (and biologically unlikely) sequence enables the chain to fold into an almost crystalline structure in a coarse-grained model, which is prevented by bond angle restrictions in the atomistic model.

**Figure 9:**
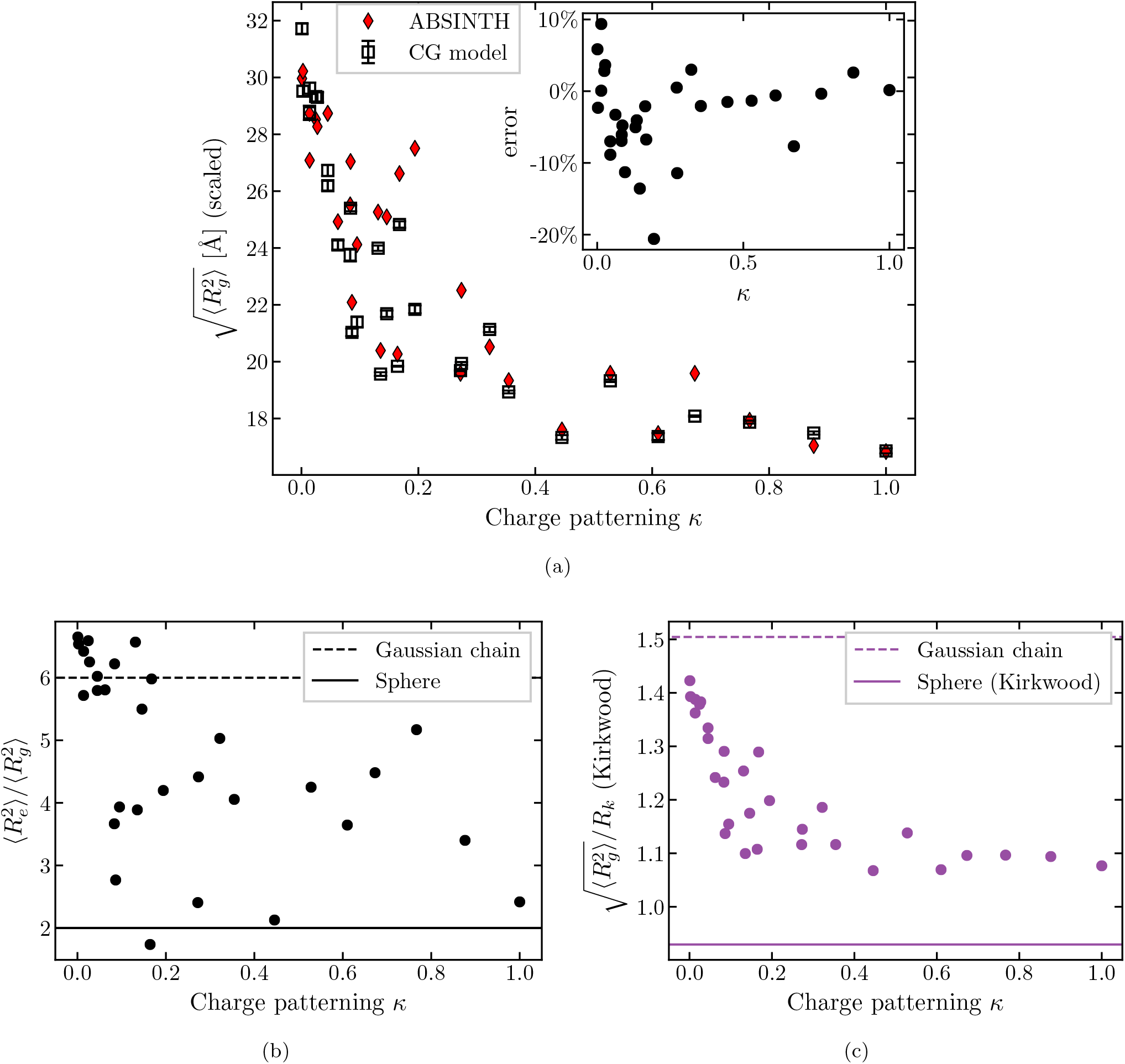
Dimensions of polymers composed of 25 positively and 25 negatively charged monomers. Same as Figure 4 but plotted with the *κ* parameter introduced by Das and Pappu [39] on the *x*-axis. *(a)* Radius of gyration compared with ABSINTH model. *(b)* Square end-to-end distance to square radius of gyration. *(c)* Radius of gyration to hydrodynamic radius (Kirkwood approximation). The dashed lines correspond to the Gaussian chain predictions, the solid lines correspond to a uniform sphere. The full sequences and their corresponding *κ* and SCD parameters are shown in Fig. 4a.

Figures 4*c* and 4*d* show the ratios of the different polymer dimensions for the different sequences. Unlike the”patchy” cohesive sequences of Figure 3, for the polyampholytic sequences the ratio of the end-to-end distance to the radius of gyration is very sensitive to the specific sequence. On the other hand, the ratio of the radius of gyration to the hydrodynamic radius is correlated with SCD and the overall compaction reflected in *R_g_*, and determines well the position of the sequence on the disorder-to-order continuum. This indicates that FRET measurements might be more indicative of local structure near the polymer ends, and cannot always used to infer the other polymer dimensions.

Notably, the smooth way in which the radius of gyration and the *R_g_*/*R_k_* ratio depend on the sequence “patchiness” (SCD) resembles the dependence of the hompolymer dimensions on the cohesiveness parameter *ϵ*. Moreover, it has been shown [58] that SCD and *R_g_* are both correlated with the critical temperature of the IDP phase separation, establishing a connection between the SCD and the mean field Flory parameter *χ* that describes the average attraction between chain monomers [15, 84]. Thus, the effect of changing the “patchiness” of a polyampholite sequence (quantified via SCD) on the radius of gyration and the phase separation behaviour of IDPs is analogous to adjusting the global average cohesiveness of the polymer. Thus, each polyampholite sequence can be mapped onto an effective homopolymer model, by finding the homopolymer *ϵ* that produces the same *R_g_*/*R_k_* ratio as the heterogeneous sequence, as shown in Figure 4d and Figure 5b.

**Figure 5:**
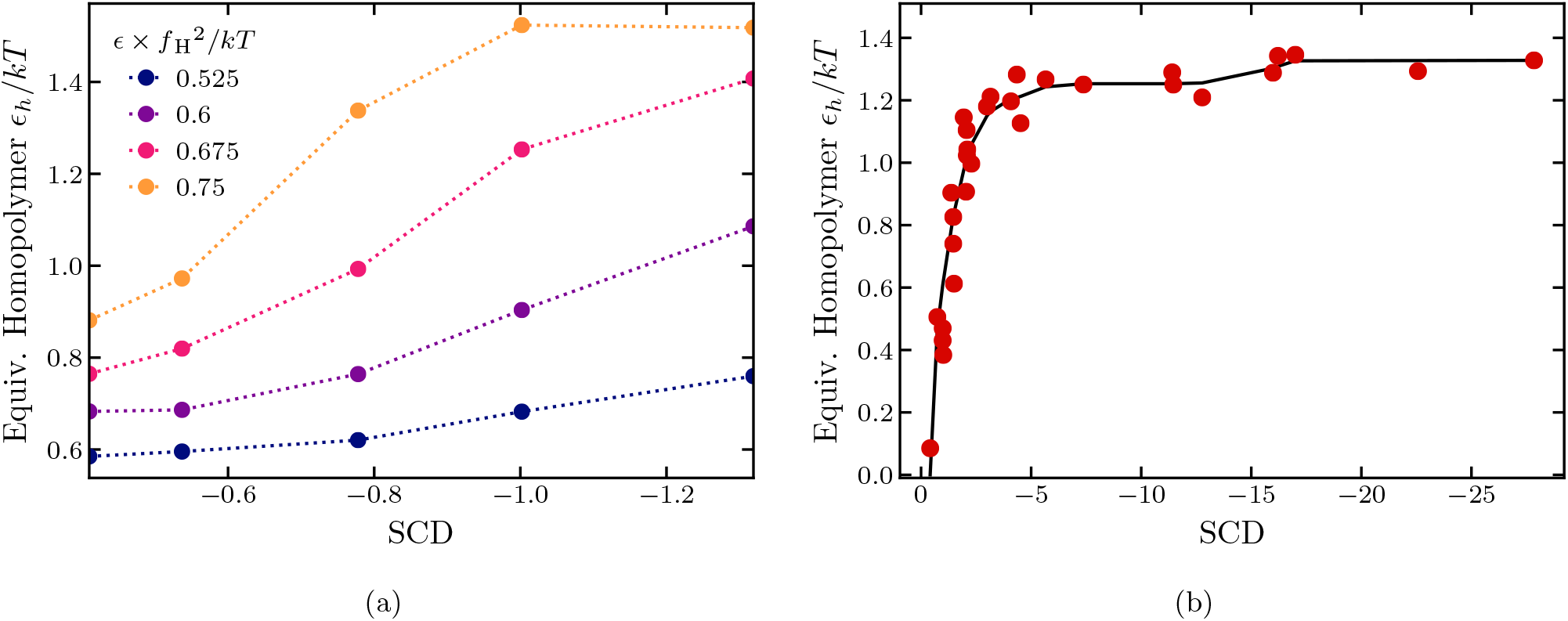
Equivalent Homopolymer model. *(a)* Cohesiveness *ϵ_h_* of the effective homopolymer model that reproduces the radii of gyrations of sequences with cohesiveness *ϵ* shown in Figure 3 and Table 1, as a function of their SCD. *(b)* Cohesiveness *ϵ_h_* of the effective homopolymer model that reproduces the *R_g_*/*R_k_* ratio of the sequences composed of 25 positively and 25 negatively charged monomers shown in Figure 4, as a function of their SCD value. The red dots show the individual correspondence for each sequence based on Figure 3. The black line is the smoothed isotonic regression *R_g_*/*R_k_* vs. SCD; see text.

Similar mapping can be achieved for the “HP” sequence consisting of a mixture of neutral and cohesive monomers that has been discussed above, as shown in Figure 5a and Figure 3b. For moderate values of inter-monomers cohesiveness e, the effective cohesiveness of a corresponding homopolymer, *ϵ_h_*, is well approximated by *ϵ_h_* = (*e*^*a*(*ϵ*)^ – 1)/*b*(*ϵ*), where *a* and *b* are fitting parameters.

### 3.2 Dynamics of IDP conformational reconfiguration

Fluctuations in the distance between the donor and the acceptor fluorophores, usually placed at the ends of the chain, result in fluctuations of the fluorescence intensity. Correlations in the fluorescence intensity fluctuations, measured through the combination of FRET and fluorescence correlation spectroscopy (FCS) provide information about the internal dynamics of the chain [7, 41]. The outcomes of such experiments have generated several puzzling results, and are still incompletely understood. In particular, increase in the denaturant concentration that causes swelling of the end-to-end distance, has been observed to correspond to the decrease in the end-to-end distance reconfiguration time, contrary to the naive expectation that the reconfiguration time would increase with the longer end-to-end distances [41, 85]. These observations can potentially be attributed to the “internal friction” resulting from several intra-chain interactions at lower denaturant concentrations, but the physical and molecular origin of internal friction in IDPs is still under debate [7, 42, 43]. Theoretical approaches based on Rouse (and Zimm) like models can capture some of the experimentally observed effects but often rely on *ad hoc* assumptions about the end-to-end probability distribution [86–88].

In this section, motivated by the experimental studies of the dynamics of the IDP configurational changes [41, 85, 89, 90], we investigate the dynamics of the end-to-end distance of IDPs using several coarse-grained examples. We focus on the dynamics of the two experimentally motivated quantities: the auto-correlation times of the end-to-end-vector and the end-to-end distance.

The normalized auto-correlation function of the end-to-end vector is defined as:

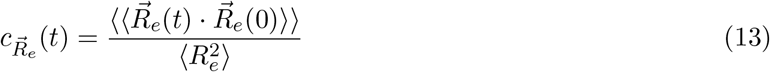

The double angle brackets represent averaging over both the initial conditions and realizations of the random simulation trajectories. The decay time of this function is referred to as the “relaxation time” of the end-to-end vector or the “rotation time” [23, 86].

The normalized auto-correlation function of the end-to-end distance is defined as:

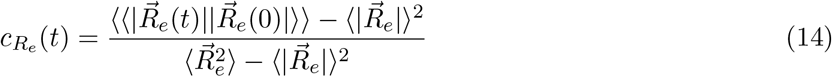

The decay time of this function is referred to as the the “reconfiguration” time. It excludes contributions from the rotation modes of the entire polymer, and is closer to the reconfiguration times captured by FRET and FCS experiments [7, 86, 87].

We calculate the correlation times *τ* of the end-to-end vector and the end-to-end distance as the integral of their normalized auto-correlation functions: 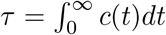 where *c*(*t*) is 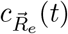 or *c_R_e__*(*t*) [82]. For computational convenience, the upper limit of the integral was cut off at *t* = 3*τ_e_* where *τ_e_* satisfies *c*(*τ_e_*) = *e*^−1^. Other methods, such as approximating the auto-correlation by an exponentially decaying function, produce substantially the same results, although further investigation is required to understand the shapes of the auto-correlation functions [82, 91].

To understand the effects of sequence composition and patterning, we focus on four sequences composed of cohesive (“H”) and neutral (“P”) monomers comprising *N* = 100 monomers each. The first sequence is the homopolymer introduced in Section 3.1.1, “(H)_100_”. The second sequence consists of a repeated “HP” motif, “(HP)_50_”. The two remaining sequences consist of a repeated “HPP” motif: one with cohesive monomers at the ends, “(HPP)_33_*H*” and the other with neutral monomers at the ends, “P(HPP)_33_”.

For the homopolymer, the cohesive interactions strengths ranged from 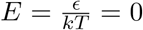 to 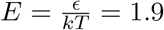 inclusive, in intervals of 0.1. Because different heteropolymer sequences have different fractions of cohesive monomers, in order to compare end-to-end dynamics for comparable chain dimensions for the “(HP)_50_” sequence the cohesive interaction strengths were: 0.5, 1, 1.5, 2.0, 2.2, 2.4, 2.6, 2.8, 3, 3.2, 3.4, 3.6, 3.8, 4, 4.2, 4.4, 4.6, and 4.8; for the “(HPP)_33_H” and “P(HPP)_33_” sequences, the cohesive interaction strengths were: 0.5, 1, 1.5, 2, 2.5, 3, 3.5, 4, 4.5, 5, 5.1, 5.2, 5.3, 5.4, 5.5, 5.6, 5.7, 5.8, 5.9, and 6.

For each 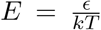, 240 runs were performed, each lasting ~ 1.8 × 10^7^ steps, with a time step of Δ*T* = 0.001 in simulation units. Each run began with a self avoiding walk initial condition. The first 2 × 10^6^ steps were excluded from the analysis, and averages were taken both over the time and the ensemble. For each run, the auto-correlation functions were calculated using the Fast Correlation Algorithm [91]. The auto-correlation functions were subsequently averaged over different runs for each *ϵ*. These auto-correlation functions are shown in Figure 6.

**Figure 6:**
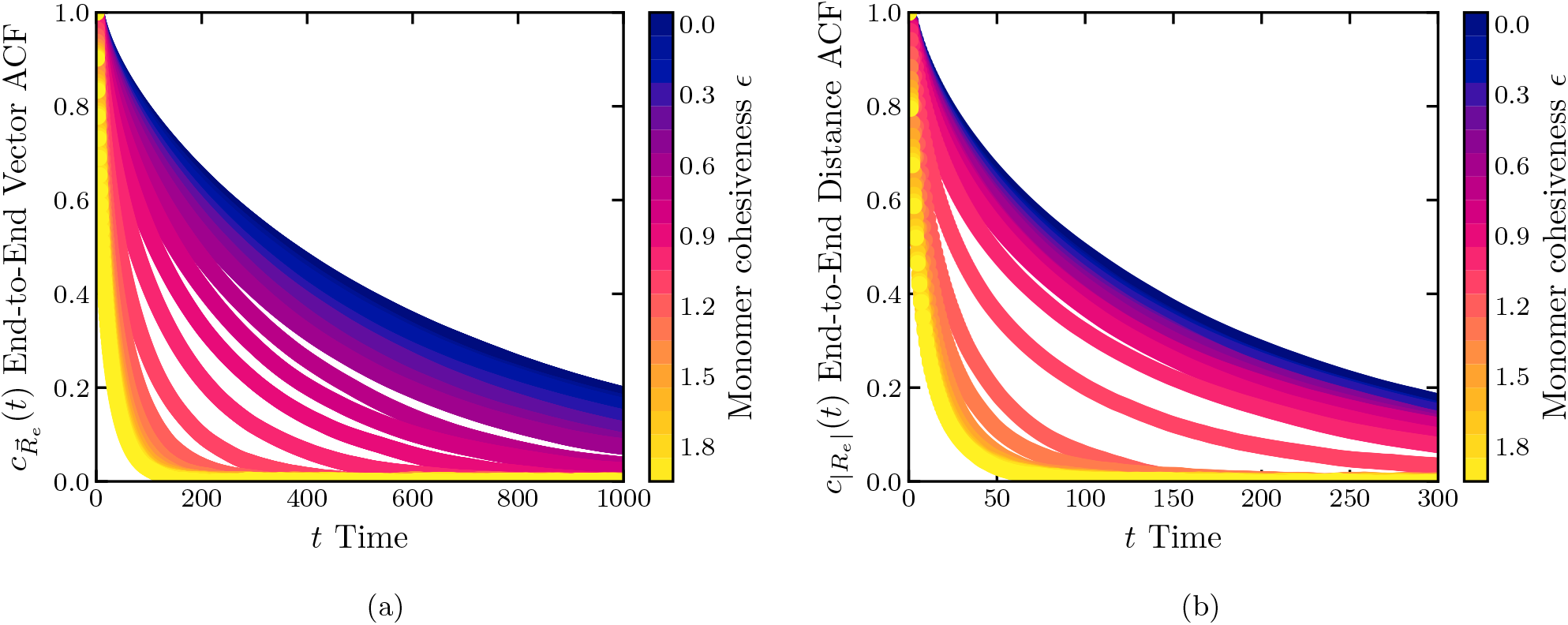
Illustrative normalized autocorrelation functions (ACF) of the *(a)* end-to-end vector and *(b)* and end-to-end distance. Results are for the homopolymer model with *N* = 100.

For all four sequences, the end-to-end vector relaxation time decreases monotonically with *ϵ*, 7*a*. As expected, in the swollen regime, above the *θ*-point the end-to-end relaxation rotation time is well described by the classical Zimm time in good and *θ*-solvent regimes 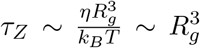, shown by the black line [23]. More globular chains below the *θ*-point start deviating from the Zimm time, although the agreement is still good for all sequences except “(HPP)_33_*H*”. We return to the special behavior of this sequence below.

The behavior of the end-to-end distance reconfiguration time is shown in Figure 7*b*. Similar to the relaxation time of the end-to-end vector, the reconfiguration time decreases monotonically with the chain compactness for the homopolymer, the “(HP)_50_”, and the “P(HPP)_33_” sequences, although the dependence does not obey the Zimm law anymore. By contrast, for the “(HPP)_33_H” sequence that has cohesive monomers at the ends, the reconfiguration time is a non-monotonic function of the chain dimensions in the compact regime below the *θ*-point.

**Figure 7:**
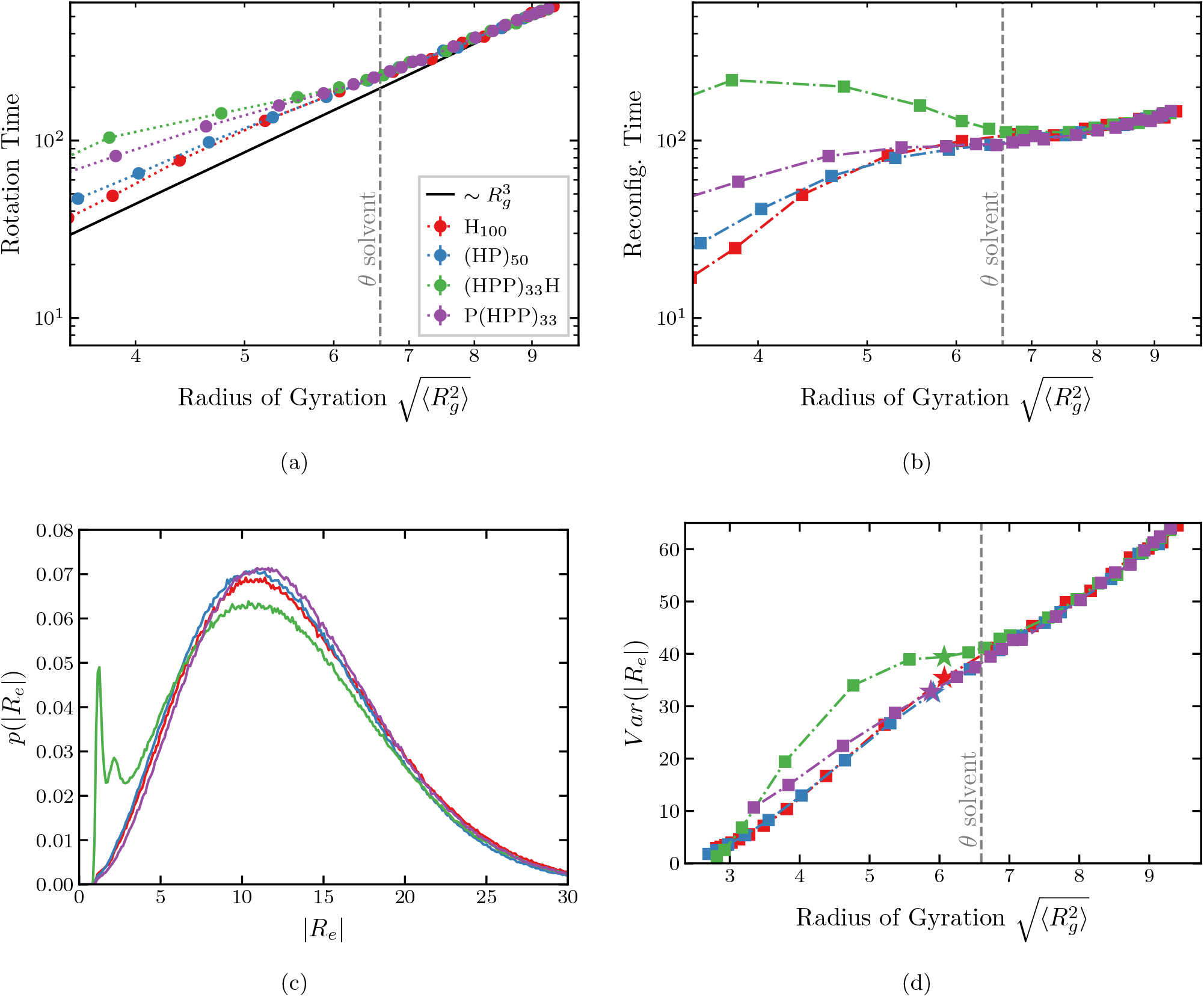
Relaxation times of the end-to-end vector and distance. *(a)* Relaxation time of the end-to-end vector (“rotation time”) and *(b)* the end-to-end distance (“reconfiguration time”) for the different sequences indicated in the legend of *(a)*. The *x*-axis shows the mean square radius of gyration controlled by monomer cohesiveness in the simulations. *(c)* End-to-end distance probability distribution for chains with different sequences. Red line: H_100_ sequence; *ϵ*/*kT* = 0.9. Blue line: (HP)_50_ sequence; *ϵ*/*kT* = 3.2. Green line: (HPP)_33_H sequence; *ϵ*/*kT* = 5.4. Purple line: P(HPP)_33_; *ϵ*/*kT* = 5.6. The radius of gyration *R_g_* ≈ 6 ± 0.1 for all sequences (see *(d)*). *(d)* Variance of the end-to-end distance as a function the radius of gyration of the chains. Stars indicate the radii of gyration of the sequences for the parameter values in *(c)*. Deviation of the green line from the others below the *θ*-point reflect the emergence of the secondary peak in the end-to-end distribution in *(c)*. See text.

This behavior can be understood by examinining the distributions of the end-to-end distances for the chains of different sequences (Fig. 7*c*). For the homompolymer, the “(HP)_50_”, and the”P(HPP)_33_” sequences, the end-to-end distance distributions have a single peak around a typical value of the end-to-end-distance. However, for the “(HPP)_33_H” sequence, additional peaks emerge immediately after the polymer compacts beyond the *θ* solvent condition. This feature is further illustrated in Figure 7*d* which shows the variances of the end-to-end distance distributions for the four sequences as a function of the compactness. Above the *θ*-point, the variances are identical for all sequences. By contrast, below the θ-point For the “(HPP)_33_H” chain with cohesive monomers the variance is significantly higher than fr the other sequences, reflecting the emergence of the secondary compact conformation shown in Fig. 7*c*).

The transition between these two conformations - with ends bound to each other and far apart, respectively - is responsible for the non-monotonic dependence of the reconfiguration time on the chain compaction exhibited in Fig. 7*b*). Namely, for the “(HPP)_33_H” sequence the free energy landscape in conformation space is more rugged, and the polymer is sampling a few highly probable conformations rather than smoothly transitioning between conformations of a Gaussian chain. In conclusion, the anomalous behavior of the reconfiguration time arises from the particular properties of the sequence.

These results have important implications for the interpretation of FRET and FCS experimental results that commonly assume a Gaussian end-to-end distribution, and where the interaction between the FRET dyes can be or importance [90]. This effect might explain the behaviour observed for chemically denatured proteins and IPDs in FRET and FCS experiments [41, 85, 89, 90].

## 4 Summary and Discussion

The absence of agreed upon computational models of IDPs makes the iterpretation of the experimental results difficult, and often leads to apparent discrepancies. Although specific models have been successful in explaining experimental results in a number of systems, the full picture of the effects of amino acid composition and sequence specifity on the behavior of IDPs and IDRs still remains incomplete. In this paper, we have systematically investigated the effects of internal interactions and sequence heterogeneity on the dimensions of IDPs, with applications to several experimental techniques. Although we use a minimal coarse-grained model, our results are likely to be general, as illustrated by their excellent agreement with the results obtained using atomistic simulations.

For the homopolymer model with internal cohesiveness, which serves as a ”null model” against which the more complex models can be benchmarked, increase in the cohesiveness results in a consistent compaction of all the polymer dimensions (end-to-end distance *R_e_*, radius of gyration *R_g_*, and the hydrodynamic radius *R_h_*). The degree of compaction differs for each of the polymer dimensions and their ratios (*R_e_*/*R_g_* and *R_g_*/*R_k_*). We also found that the conformations of the homopolymer are aspherical for low values of cohesiveness, and the ratio of end-to-end distance to the radius of gyration is correlated with asphericity. In terms of dynamical quantities, both the rotation and the reconfiguration times of the end-to-end distance decreases monotonically with the polymer compactness caused by the increase in the cohesiveness.

Sequence heterogeneity can significantly modulate the polymer dimensions independently of the composition or the attraction strength between cohesive monomers. For polymers composed of mixtures of cohesive and neutral monomers, an increase in the size of cohesive “patches” resulted in the more significant compaction of the polymer, reflected in all dimensions and their ratios. Nevertheless, the overall behavior of these polymers can be semi-quantitatively mapped onto that of a simple homopolymer with an appropriately chosen value of the average cohesiveness. For low values of cohesiveness below the *theta*-point, this effective cohesiveness is simply proportional to the square of the fraction of the cohesive monomers in the chain, reflecting the mean field reduction in the average number of inter-monomer contacts. For more cohesive sequences in a compact regime, the mean field description starts to break down, and the effective homopolymer cohesiveness depends on the “patch” size. In this regime, the effective cohesiveness correlates with the SCD of the sequence, which also was shown to correlate with the macroscopic Flory parameter describing the mean field cohesive behavior of single chains, and their collective properties such as the phase separation.

Presence of monomers of both positive and negative charges in the sequence can have dramatic effect on polymer dimensions, as described in Section 3.1.1. Notably, in this case the mean field description completely breaks down due to the cancellation of interactions between oppositely charged monomers. To study the effects of charge patterning, and to validate our model, we studied a polyampholyte sequence composed of positively and negatively monomers. The dimensions of the polyampholytes predicted by our coarse-grained model were similar to those predicted by an all-atom model with explicit ions, and exhibited the same trends. Overall, the radius of gyration, *R_g_* and ratio of the radius of gyration to the hydrodynamic radius, *R_g_*/*R_k_*, monotonically decayed with the sequence patterning parameters SCD and *κ*, enabling mapping from SCD onto an average cohesiveness of an effective homopolymer model. These results are consistent with the findings that the SCD correlates with the phase transition temperature and thus with the Flory parameter *χ*.

However, unlike for the cohesive/neutral chains, for the polyampholytes the end-to-end distance *R_e_* and the ratio *R_e_*/*R_g_* were highly sequence specific. This partial decoupling between the *R_e_* and *R_g_*, arising from the high sensitivity of *R_e_* to the details of the sequence at the chain ends, is in agreement with previous observations and modeling. Thus, while *R_g_*/*R_k_* ratio appears to be a robust parameter that locates the IDP on the order-disorder continuum and is useful in the interpretation of experiments, the end-to-end distance *R_e_* and its ratio *R_e_*/*R_g_* are not, and care should be exercised while interpreting FRET experiments.

Nevertheless, rather than being the source of a discrepancy, the combined measurements of several polymer dimensions can guide the interpretation of experimental results and the inference of the internal interactions of an IDP. For example, the ratio between radius of gyration and hydrodynamic radius can reveal the location of a particular IDP on the disorder-to-order continuum, while the ratio of end-to-end distance to radius of gyration may reveal the relative importance of the direct end-to-end interactions.

The sensitivity of the end-to-end distance to the properties of the monomers at the chain ends shows itself also in the end-to-end dynamics. Puzzlingly, both IDPs and chemically denatured proteins can exhibit a non-monotonic dependence of the end-to-end distance reconfiguration times on denaturant concentrations and the associated chain compaction. Molecular dynamics studies have proposed “internal friction” as the source of this behaviour, but its microscopic origin still remains unclear, and the reconfiguration dynamics is still not fully understood.

The coarse grained model of this paper shows that the end-to-end distance distribution and thus the end-to-end distance reconfiguration time is sensitive to the properties of the monomers near the chain ends. Sequence with cohesive monomers at the ends exhibited a regime where the reconfiguration time increases with the compaction of the polymer dimensions, qualitatively following the experimental observations. This increase was contingent on the emergence of multiple peaks in the end-to-end distance distribution due to the presence of cohesive monomers at the ends, indicating bi-stability between a compact and a swollen conformations. Chains with more homogeneous sequences explore Gaussian conformational landscapes and have faster end-to-end distance reconfiguration times, while those with more heterogeneous sequences explore more distant conformational states and therefore have slower re-configuration times. This difference between the conformational ensembles would not appear in a static measurement of polymer dimensions. This emphasizes again the importance of sequence for the end-to-end dynamics and statics, and might contribute to the understanding of the origin of the “internal friction” of IDPs.

Our coarse grained model offers a powerfull tool for the interpretation of the equilibrium and dynamics experiments without resorting to all atom simulations. The coarse-grained model encapulates a wide range of general IDP behaviors, semi-quantitatively agrees with atomistic simulations, and serves as the baseline for mode complex models. Future investigation will apply the coarse-grained model to specific cases of IDPs to understand their behavior in multi-chain assemblies and their interaction and binding with other proteins.

## A Appendix

### A.1 Hydrodynamic Radius and the Kirkwood Approximation

The hydrodynamic radius *R_h_* is the radius of a solid sphere that has the same diffusion coefficient as the polymer chain. It depends not only on the equilibrium properties of the conformational ensemble but also on the dynamical intra-chain correlations mediated via fluid flow around and within the chain. The diffusion coefficient of a polymer chain is inversely proportional to the hydrodynamic radius via the Stokes-Einstein relation [22, 23]:

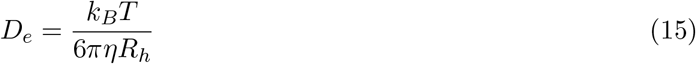

where *k_B_* is Boltzmann’s constant, *T* is the absolute temperature, *η* is the solvent viscosity. *D_e_* can be calculated from displacement of the polymer’s centre of mass as [59, 61]::

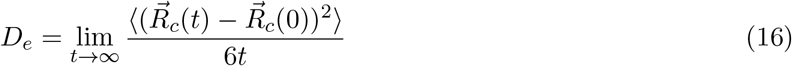

The average is over trajectories and the initial conditions.

Kirkwood and Riseman [92] introduced a pre-averaging approximation for the hydrodynamic interactions between the monomers, allowing to calculate the approximate hydrodynamic radius from just the equilibrum ensemble of conformations. The Kirkwood approximation for the diffusion coefficient of a polymer, using the Oseen tensor for hydrodynamic interactions, is [59]:

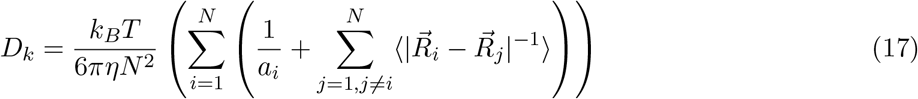

Here, *a_i_* is the hydrodynamic radius of a monomer *i*, and the average is over the equilibrium ensemble of conformations. The inverse of the approximation to the hydrodynamic radius is defined as:

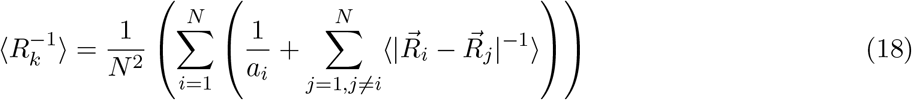

Using Brownian dynamics simulations with implicit hydrodynamic interactions, Liu et al. [59] and Schmidt et al. [93] have previously found that the Kirkwood approximation overestimates the hydrodynamic radius by < 4% for a SAW and a worm-like chain model. In this paper, we extended the comparison between the Kirkwood approximation and the hydrodynamic radius to all values of cohesiveness. The Kirkwood approximation overestimates the true diffusion coefficient by 3-5% in agreement with other studies [59, 93]. In the poor solvent regime the relative difference increases to beyond 10% and is larger for longer polymers (Fig. 1d).

### A.2 Scaling Exponent of Radius of Gyration

We investigated the effects of monomer cohesiveness *ϵ* on the scaling exponent *ν* of the radius of gyration of a homopolymer with the number of bonds 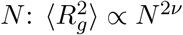.

We performed simulations of homopolymers with *N* + 1 = 50,100,150, 200, 300, 400 monomers at *ϵ* = 0,0.64,0.7,0.75 kT. All other simulation parameters were the same as the homopolymer model of Section 2. The total runtime (number of steps) and number of independent runs (from different initial conditions) varied. Simulations where initialized from a self-avoiding walk (*N* + 1 = 50,100,150,200) or a random walk initial condition (*N* + 1 = 200,300, 400). Correlation functions of 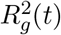 were calculated for *N* + 1 = 50,100,150 and fit with exponential decays in order to estimate the correlation times *τ_N_*. The initial 2*τ_N_* or more of each simulation were excluded from the analysis. The error bars were estimated as 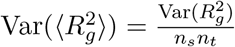 where *n_s_* is the number of independent runs and 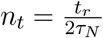, where *t_r_* is the minimum simulation time included in the analysis for that *N* [94].

Figure 8 shows the dependence of the radius of gyration on the number of bonds *N* for different monomer cohesiveness *ϵ*; For *ϵ* = 0, as expected, the chain behaves as a self-avoiding random walk in a god solvent with *ν* = 0.588 ± 0.001 [23]. The location of the *θ*-point, where *ν* = 0.5, is located between *ϵ* = 0.74 kT and 0.75 kT. This is consistent with the second virial coefficient for the inter-monomer interaction being 0 at *ϵ* ≈ 0.64 kT. The exact location of the *θ*-point depends on the details of the repulsive and attractive potentials used in the model [70].

**Figure 8:**
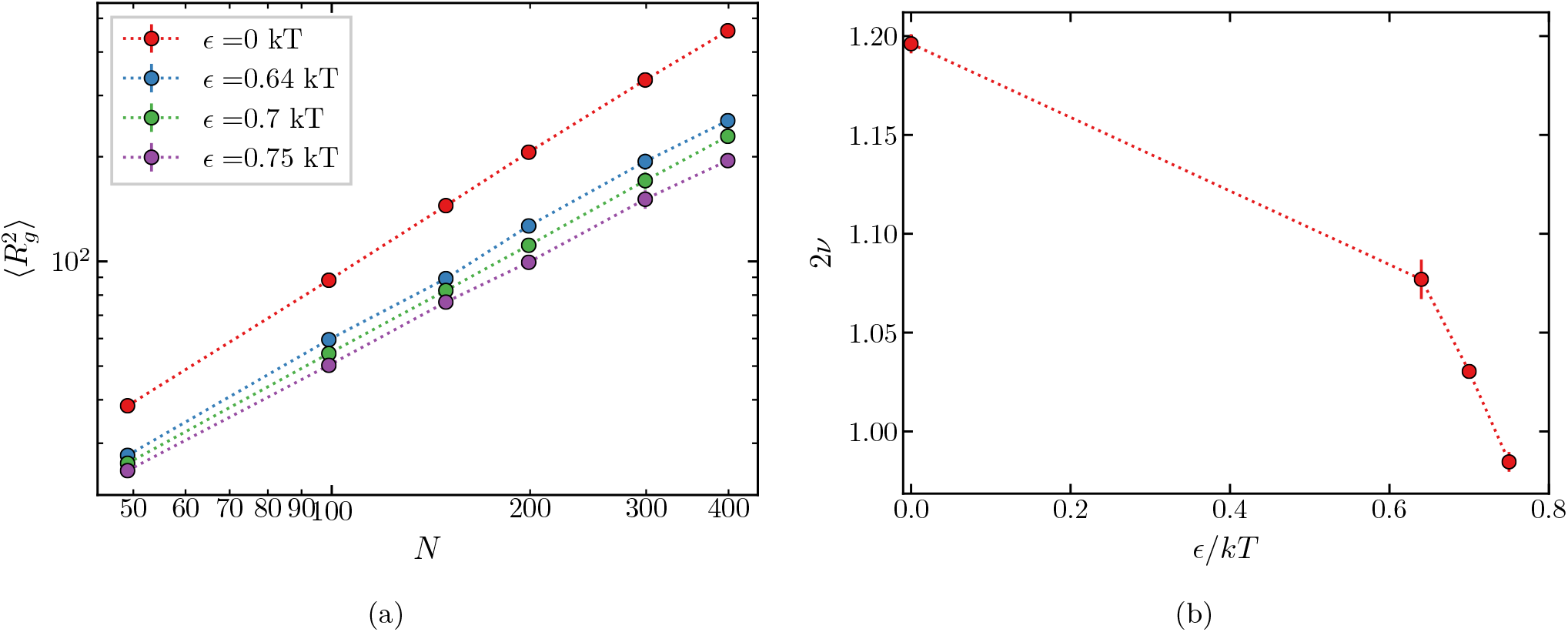
(a) Dependence of radius of gyration on the number of bonds in the chain for different monomer cohesiveness *ϵ*. (b) The variation of the scaling exponent of 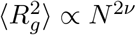 with monomer cohesiveness *ϵ*.

### A.3 Comparison between sequence patterning parameter SCD and *κ*

Figure 9 shows the polymer dimensions and the ratios of dimensions of polyampholyte sequences plotted against the charge sequence parameter *κ* introduced by Das and Pappu [39].

